# The balancing act of *Nipponites mirabilis* (Nostoceratidae, Ammonoidea): managing hydrostatics during a complex ontogenetic trajectory

**DOI:** 10.1101/2020.06.11.145813

**Authors:** David J. Peterman, Tomoyuki Mikami, Shinya Inoue

## Abstract

*Nipponites* is a heteromorph ammonoid with a complex and unique morphology that obscures its mode of life and ethology. The seemingly aberrant shell of this Late Cretaceous nostoceratid seems deleterious. However, hydrostatic simulations suggest that this morphology confers several advantages for exploiting a quasi-planktic mode of life. Virtual, 3D models of *Nipponites mirabilis* were used to compute various hydrostatic properties through 14 ontogenetic stages. At each stage, *Nipponites* had the capacity for neutral buoyancy and was not restricted to the seafloor. Throughout ontogeny, horizontally facing to upwardly facing soft body orientations were preferred. These orientations were aided by the obliquity of the shell’s ribs, which were parallel to former positions of the aperture during life. Static orientations were somewhat fixed, inferred by stability values that are slightly higher than extant *Nautilus*. The initial open-whorled, planispiral phase is well suited to horizontal backwards movement with little rocking. *Nipponites* then deviates from this coiling pattern with a series of alternating U-shaped bends in the shell. This modification allows for proficient rotation about the vertical axis, while possibly maintaining the option for horizontal backwards movement by redirecting its hyponome. These particular hydrostatic properties likely result in a tradeoff between hydrodynamic streamlining, suggesting that *Nipponites* assumed a low energy lifestyle of slowly pirouetting in search for planktic prey. Each computed hydrostatic property influences the others in some way, suggesting that *Nipponites* maintained a delicate hydrostatic balancing act throughout its ontogeny in order to facilitate this mode of life.

## Introduction

Heteromorph ammonoids are ectocochleate cephalopods whose shells undergo changes in coiling throughout ontogeny. The seemingly aberrant shape of some heteromorph ammonoids piques curiosity about their enigmatic modes of life and life habit. Arguably, the most bizarre and conspicuous of all heteromorph genera is the Late Cretaceous (Turonian – Coniacian) nostoceratid, *Nipponites* (Fig 1). Previous research has largely focused on the biostratigraphic usefulness of *Nipponites* [1–5] rather than its paleobiology [6–7] and evolutionary significance [8]. The latter two areas are valuable because the morphology of this heteromorph appears deleterious to survival; seemingly defying the basic principles of natural selection [9–15]. It is more likely, however, that its functional morphology is obscured by a complex ontogenetic trajectory in shell growth. The shell of *Nipponites* is characterized by having several open planispiral (crioconic) whorls in early ontogeny, followed by a series of alternating U-bends around the earlier whorls (Fig 1); denoting some degree of regularity in coiling throughout a seemingly-aberrant ontogeny [1,16,17]. Okamoto [18–20] demonstrated that the coiling of *Nipponites mirabilis* is, in fact, well constrained and can be approximated by a few piecewise equations (alternations of sinistral and dextral helicoid phases surrounding the crioconic phase). Similarly, differential geometry has proven a useful tool in modeling these complex heteromorphs [21–22].

**Fig 1.**
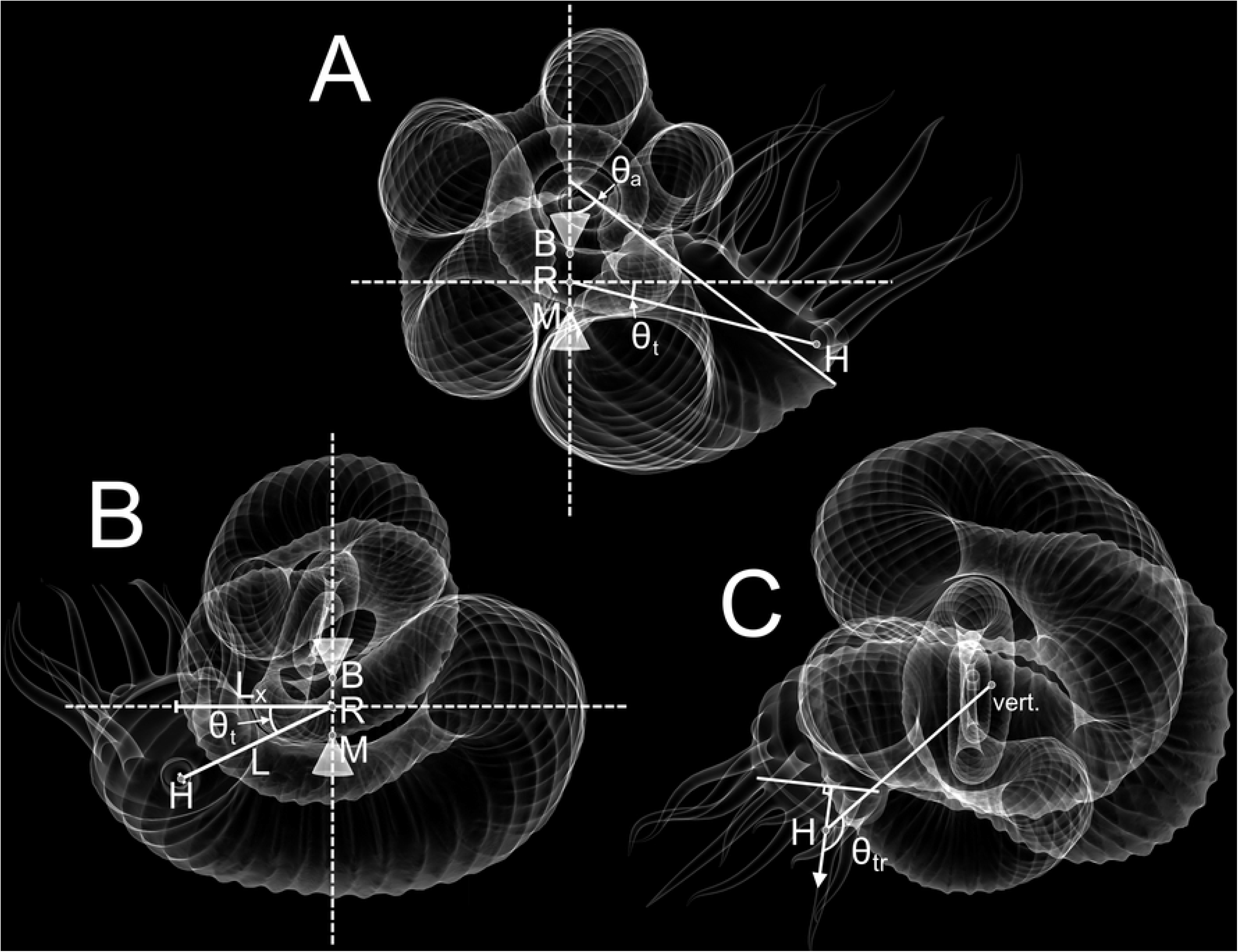
Hydrostatic Parameters of *Nipponites mirabilis*. **A**, Side view of *Nipponites* in life position showing hypothetical centers of buoyancy (B), mass (M), and the horizontal axis of rotation (R). The angle of the aperture (θ_a_) is measured as the inclination from the vertical plane. The thrust angle (θ_t_) can be used to assess the directional efficiency of movement. This angle is measured between the horizontal plane, and a line passing through R and the location of the hyponome (source of thrust; H). **B**, Front view of *Nipponites* in life position facing the aperture. This view shows the total lever arm (L) and its x-component (L_x_) which is proportionate to the amount of rotational movement about the vertical axis produced during jet propulsion. **C**, Top view of *Nipponites* in life positon showing the rotational thrust angle (θ_tr_). This angle is measured between the vertical rotation axis (vert.), which passes through B and M, and the direction of the thrust vector (arrow emanating from H). Rotational thrust angles of 90° result in idealized transmission of thrust into pure rotation.

The complex, meandering shell of *Nipponites* has invited several different interpretations regarding potential modes of life assumed by this heteromorph. The shell morphology of *Nipponites* has been compared to vermetid gastropods, and by analogy, this heteromorph has been suggested to assume a sessile and benthic mode of life [23–28]. Trueman [29] also considered *Nipponites* as a benthon, but with some degree of mobility. Other nostoceratid genera have been interpreted as negatively buoyant, benthic elements as well [28,30,31]. By similar analogy with other ‘irregularly-coiled’ mollusks, a symbiotic relationship with sponges or hydrozoans occupying the free space between the whorls of *Nipponites* has been speculated [32]; although, no fossil evidence currently supports such a relationship. Contrasting benthic interpretations, Ward & Westermann [33] suggest that *Nipponites occidentalis* was capable of a planktic mode of life based on approximate calculations of organismal density. This mode of life is supported by Okamoto [19] for *Nipponites mirabilis* due to the oscillation of rib obliquity of the shell. Changes in rib obliquity suggests that some proper orientation of the soft body was preferred, which would not matter during a negatively buoyant condition. Favoring a planktic mode of life, Westermann [6] inferred *Nipponites* was an occupant of the epipelagic, oceanic waters, perhaps as a vertical migrant or planktic drifter. This morphology is certainly not streamlined, suggesting that it would have experienced considerably more hydrodynamic drag than its planispiral counterparts. The unique shell of this genus raises questions regarding how its changes in coiling may reflect the modification of syn vivo hydrostatic properties; a tactic observed in other morphotypes of heteromorph ammonoids [17,19,20,34-39].

### Hydrostatic properties of heteromorph ammonoids

The ability of ectocochleate cephalopods to attain neutral buoyancy is fundamental to reconstruct their modes of life. The variable interpretations for nostoceratid modes of life illustrate the importance of new techniques to determine the physical properties that would have acted on these living cephalopods. A neutrally buoyant condition is achieved when the total organismal mass is equal to the mass of the water displaced by the living animal. This depends upon the body chamber to phragmocone ratio. If the phragmocone (the chambered portion of the shell) is too small, the living cephalopod would not be able to compensate for its organismal weight and it would become negatively buoyant [34,36,40]. This condition also depends upon shell thickness and the densities assigned to each component of the living animal, which have been somewhat variable in previous research [39,41].

Previous studies have demonstrated that heteromorph ammonoids may have been able to achieve much different life orientations than their planispiral counterparts [20,29,34–39,42–45]. These living cephalopods would have assumed some static orientation when their centers of buoyancy and mass were vertically aligned [41,46,47] (Fig 1). The difficulty to which these living cephalopods could deviate from their static orientation depends on hydrostatic stability, which is proportionate to the separation between the centers of buoyancy and mass [20]. High stability would have reduced the influence of external forms of energy on orientation, but would have simultaneously made it more difficult for the living cephalopod to self-modify its orientation [36].

The directional efficiency of movement (thrust angle) depends upon the relative position of the source of thrust (the hyponome) and the center of rotation (the midpoint between the centers of buoyancy and mass; Fig 1A, B). Thrust energy produced by jet propulsion is more efficiently transmitted into movement in the direction where the hyponome and center of rotation are aligned [20,38,39,48,49]. If these two points were horizontally aligned (thrust angle of zero), more energy would be transmitted to horizontal movement with minimal rocking. The rocking behavior of extant nautilids is related to their sub-horizontal thrust angles and the retraction of the soft body during emptying of the mantle cavity [50].

A rotational component of energy is increased by turning the direction of thrust out of alignment with the centers of buoyancy and mass (the axis where idealized rotation would occur; Fig 1B, C). An increased distance of the hyponome from these two centers would therefore produce a lever arm that would impart a torque to rotate the living cephalopod about its vertical axis. This type of movement is likely to have taken place for turrilitid heteromorphs [34], as well as other morphotypes with their apertures positioned in a similar relative manner [37]. Idealized rotation about the vertical axis would occur with a long, horizontally oriented lever arm and a thrust vector adjoining its distal end with a right angle.

Each of these physical properties would have significantly constrained the hydrostatic and hydrodynamic capabilities of living *Nipponites* throughout its ontogeny. Therefore, they provide fundamental information regarding the possible modes of life and life habit for this unique ammonoid, as well as possible adaptations for locomotion and feeding.

## Methods

Virtual models were constructed to determine the syn vivo hydrostatic properties of *Nipponites mirabilis*. Construction of the shell and other model components largely follow the methods of Peterman et al. [37–39], although a CT scanned specimen was used as the base model instead of using photogrammetry (similar to the methods of Morón-Alfonso [51]). This modification from the previous methods was preferred for this species due to the complex changes in shell ornamentation (rib obliquity). These ribs are parallel to the successive positions of the aperture throughout ontogeny, therefore retaining vital information about life orientation [20]. This method for virtual reconstruction is favorable for *Nipponites* because specimens of this genus are rarely found complete; discouraging destructive sampling techniques like serial grinding tomography. Computed tomography (CT) scans of such specimens also lack contrasts of X-ray attenuation factors to distinguish the shell from its surrounding materials [52]. However, each of these tomographic techniques can provide very accurate measurements of hydrostatic properties and volumes when the specimens are adequate for imaging [52–59].

### Virtual modeling of the shell

The shell of *Nipponites mirabilis* was constructed from an initial CT scan [60] of the specimen INM-4-346 (Museum Park Ibaraki Prefectural Museum of Nature), which had a remarkable degree of preservation. Most of the ontogeny is preserved for this specimen with minimal matrix on the inside (Fig 2A). However, two portions had to be virtually reconstructed; the crushed ~5 cm section of the adoral-most body chamber, and 2) the earliest crioconic whorls that are partially embedded in a remnant of the original concretion. These two portions of the shell were reconstructed (Fig 2B) with array algorithms [37–40], which replicate a whorl section and simultaneously translate, rotate, and scale it to build the shell from the adoral direction to adapical direction (Table 1). Such arrays are similar to the morphospace parameters of Raup [61]. The CT scanned model [60] (which consists of a stack of .tiff images) was converted to the tessellated .stl format required for model reconstruction and volumetry using the program, Molcer 1.51 [62]. The external mesh of the tessellated file was isolated in order to get rid of internal features like fissures and X-ray attenuation artifacts. External defects were smoothed in Meshmixer 3.3 [63] while maintaining the curvature of neighboring, complete features. This external 3D mesh served as a stencil for the reconstruction of the missing and damaged portions of the shell. After the missing portions of the shell were combined to the model derived from the CT scan, the ornamentation was reconstructed by matching the width and amplitude of ribs with a torus shape in Blender [64], then properly oriented using the ribs present on the inner whorls. The ornamentation, reconstructed portions of the shell (Fig 2B), and the total external mesh were repaired and unified in Netfabb [65] to produce a single manifold mesh of the exterior shell. The program, Blender, was used to assign shell thickness to the external shell model based on measurements from specimen NMNS (National Museum of Nature and Science) MP35490 (Fig 3), producing a mesh denoting the entire shell without septa.

**Table 1.**
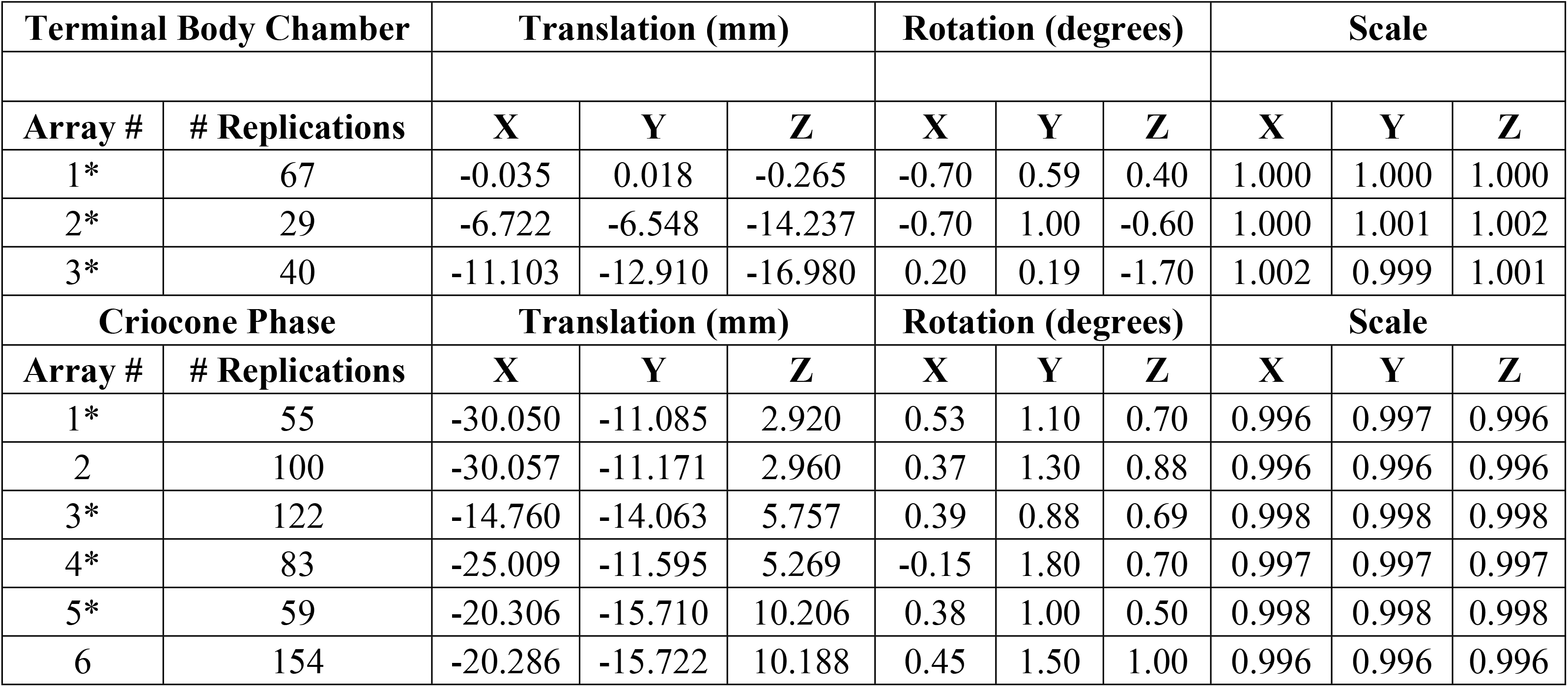
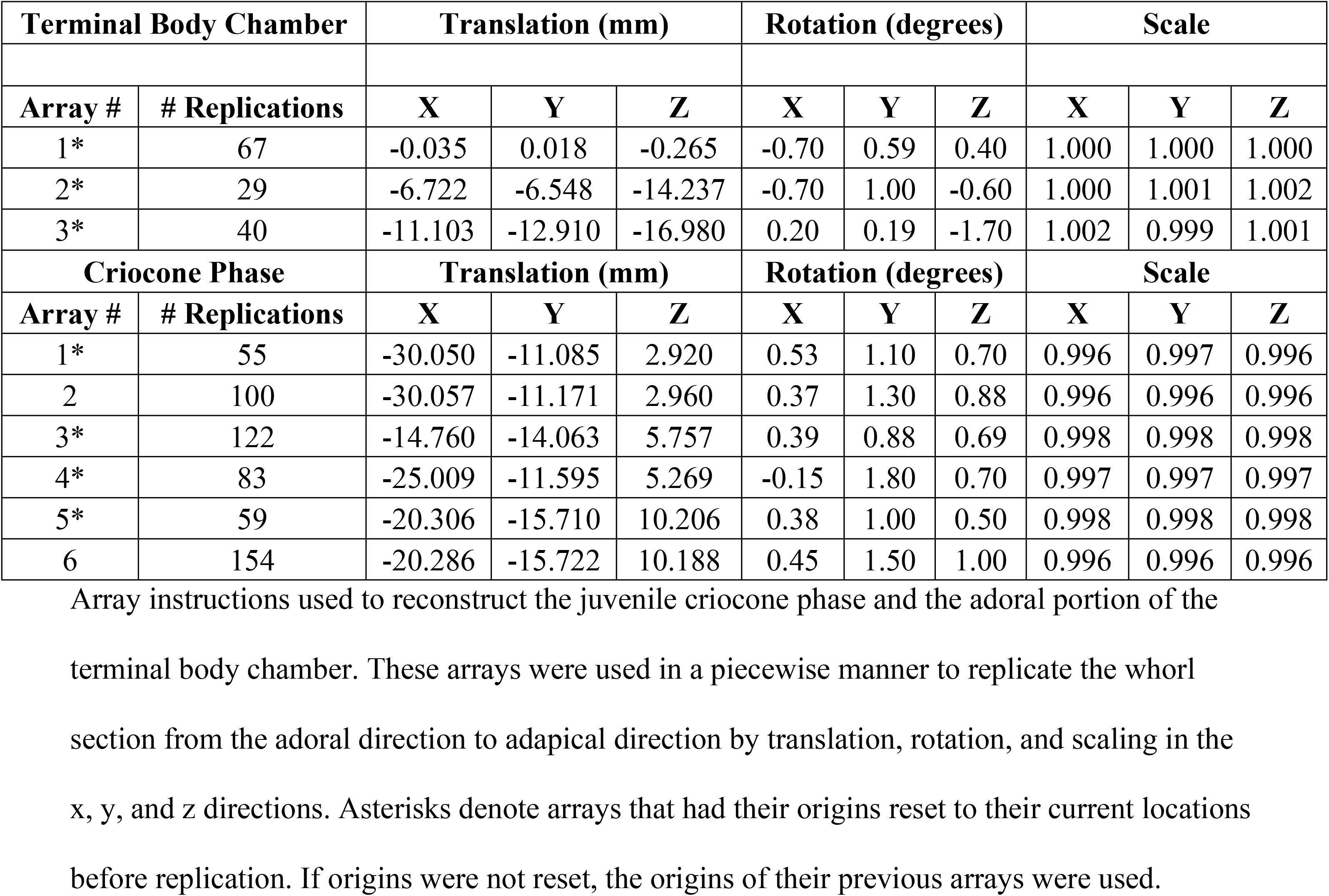
Reconstruction of the Shell

| Terminal Body Chamber |  | Translation (mm) |  |  | Rotation (degrees) |  |  | Scale |  |  |
| --- | --- | --- | --- | --- | --- | --- | --- | --- | --- | --- |
| Array # | # Replications | X | Y | Z | X | Y | Z | X | Y | Z |
| 1* | 67 | -0.035 | 0.018 | -0.265 | -0.70 | 0.59 | 0.40 | 1.000 | 1.000 | 1.000 |
| 2* | 29 | -6.722 | -6.548 | -14.237 | -0.70 | 1.00 | -0.60 | 1.000 | 1.001 | 1.002 |
| 3* | 40 | -11.103 | -12.910 | -16.980 | 0.20 | 0.19 | -1.70 | 1.002 | 0.999 | 1.001 |
| Criocone Phase |  | Translation (mm) |  |  | Rotation (degrees) |  |  | Scale |  |  |
| Array # | # Replications | X | Y | Z | X | Y | Z | X | Y | Z |
| 1* | 55 | -30.050 | -11.085 | 2.920 | 0.53 | 1.10 | 0.70 | 0.996 | 0.997 | 0.996 |
| 2 | 100 | -30.057 | -11.171 | 2.960 | 0.37 | 1.30 | 0.88 | 0.996 | 0.996 | 0.996 |
| 3* | 122 | -14.760 | -14.063 | 5.757 | 0.39 | 0.88 | 0.69 | 0.998 | 0.998 | 0.998 |
| 4* | 83 | -25.009 | -11.595 | 5.269 | -0.15 | 1.80 | 0.70 | 0.997 | 0.997 | 0.997 |
| 5* | 59 | -20.306 | -15.710 | 10.206 | 0.38 | 1.00 | 0.50 | 0.998 | 0.998 | 0.998 |
| 6 | 154 | -20.286 | -15.722 | 10.188 | 0.45 | 1.50 | 1.00 | 0.996 | 0.996 | 0.996 |

**Fig 2.**
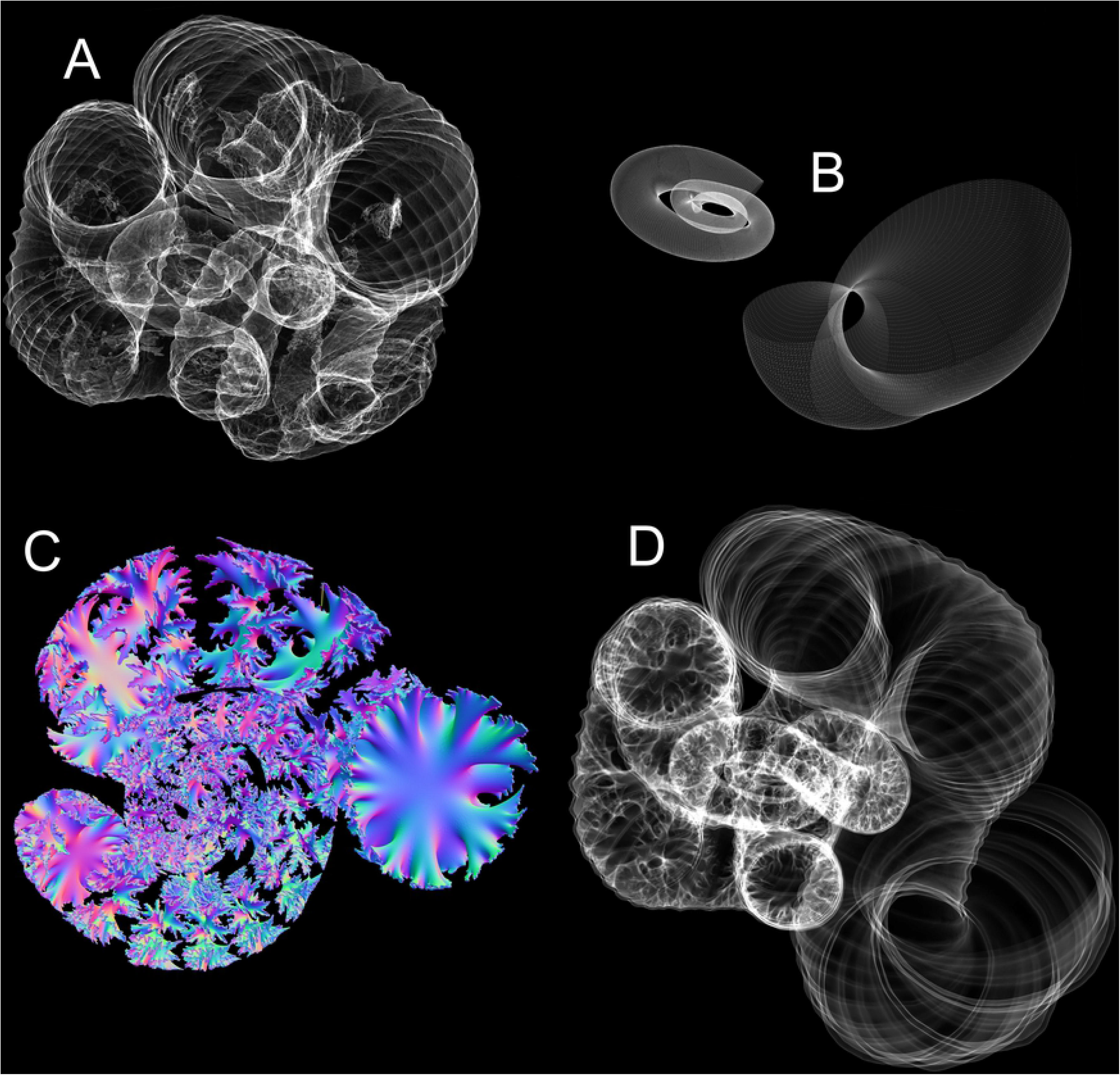
Virtual Reconstruction of the Shell of *Nipponites mirabilis*. **A**, Tessellated (.stl) 3D model generated from a CT scan [60] of specimen INM-4-346. **B**, Reconstructed adoral portion of the body chamber and inner criocone phase with arrays algorithms (Table 1). **C**, Extruded septa generated from the suture pattern. **D**, Extruded shell and septa models unified together to produce a single, manifold 3D mesh of the entire shell.

**Fig 3.**
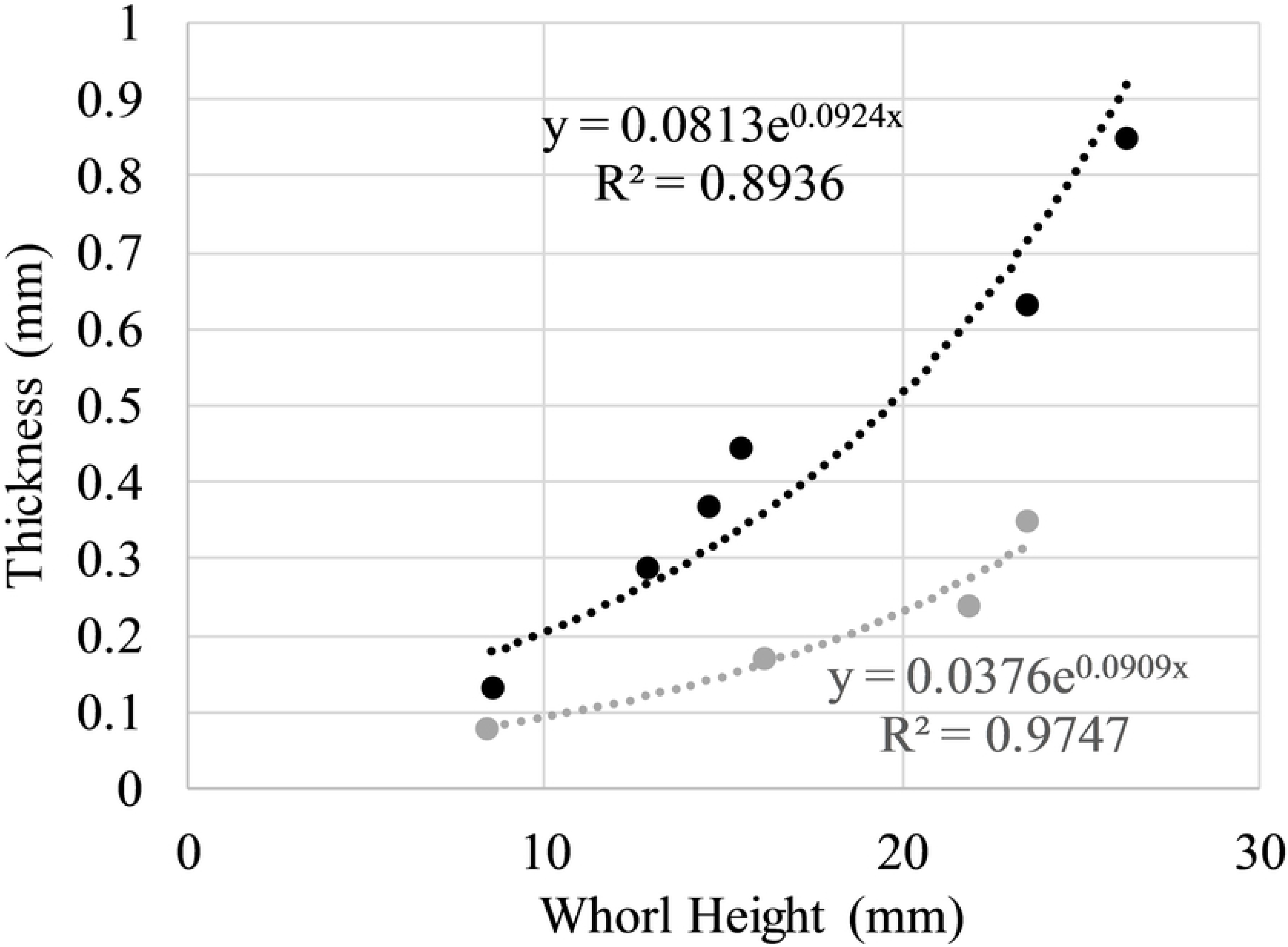
Thickness Measurements used for Virtual Model Extrusion. Thicknesses of the shell (black) and septa (grey) as a function of whorl height. Measurements were recorded from specimen NMNS PM35490 and used to define thickness in the virtual model.

Septa were constructed by recording a suture pattern from specimen NMNS PM35490 (Fig 4). The external shell of this specimen (Fig 4B, C) was removed with air abrasives and pneumatic tools under a stereoscopic microscope and the suture (Fig 4D) was recorded with a digital camera lucida. This suture was imported in the Blender workspace and the curve modifier was used to wrap it around the whorl section of the shell. This suture was then replicated and placed along the majority of the phragmocone so that adjacent lobules and folioles were almost tangential. Ontogenetic changes in the suture pattern were not considered because they probably represent only small differences in mass and its distribution. That is, each suture had the same degree of complexity and its expanded portion was placed adjacent to the venter throughout ontogeny. The crioconic, juvenile phase was reconstructed with array algorithms, which allowed septa to be duplicated with the same equations. The septa within the majority of the phragmocone were constructed by extruding the suture patterns inwards to a single point, then refining and smoothing the interior in order to approximate minimum curvature surfaces. A body chamber ratio of approximately 42% the total curvilinear length was measured from a remarkably complete specimen of *Nipponites mirabilis* from a private collection. This specimen was 3D scanned with an Artec Space Spider (to allow comparisons with the CT scanned specimen) and is housed in the morphosource database [66]. A nearly complete specimen of *Nipponites mirabilis* (MCM-A0435; Mikasa City Museum; Fig 4E) was also compared in this manner and stored in the database [66], which yielded an approximate body chamber ratio of 36%. This ratio would be slightly higher if the aperture was not partially crushed. The proper number of septa to maintain the body chamber ratio of around 42% were placed in the phragmocone and extruded based on measured thicknesses from specimen NMNS PM35490 (Fig 3). These final septa (Fig 2C) were merged with the extruded, external shell to produce a single, manifold 3D mesh of the entire, septate shell (Fig 2D).

**Fig 4.**
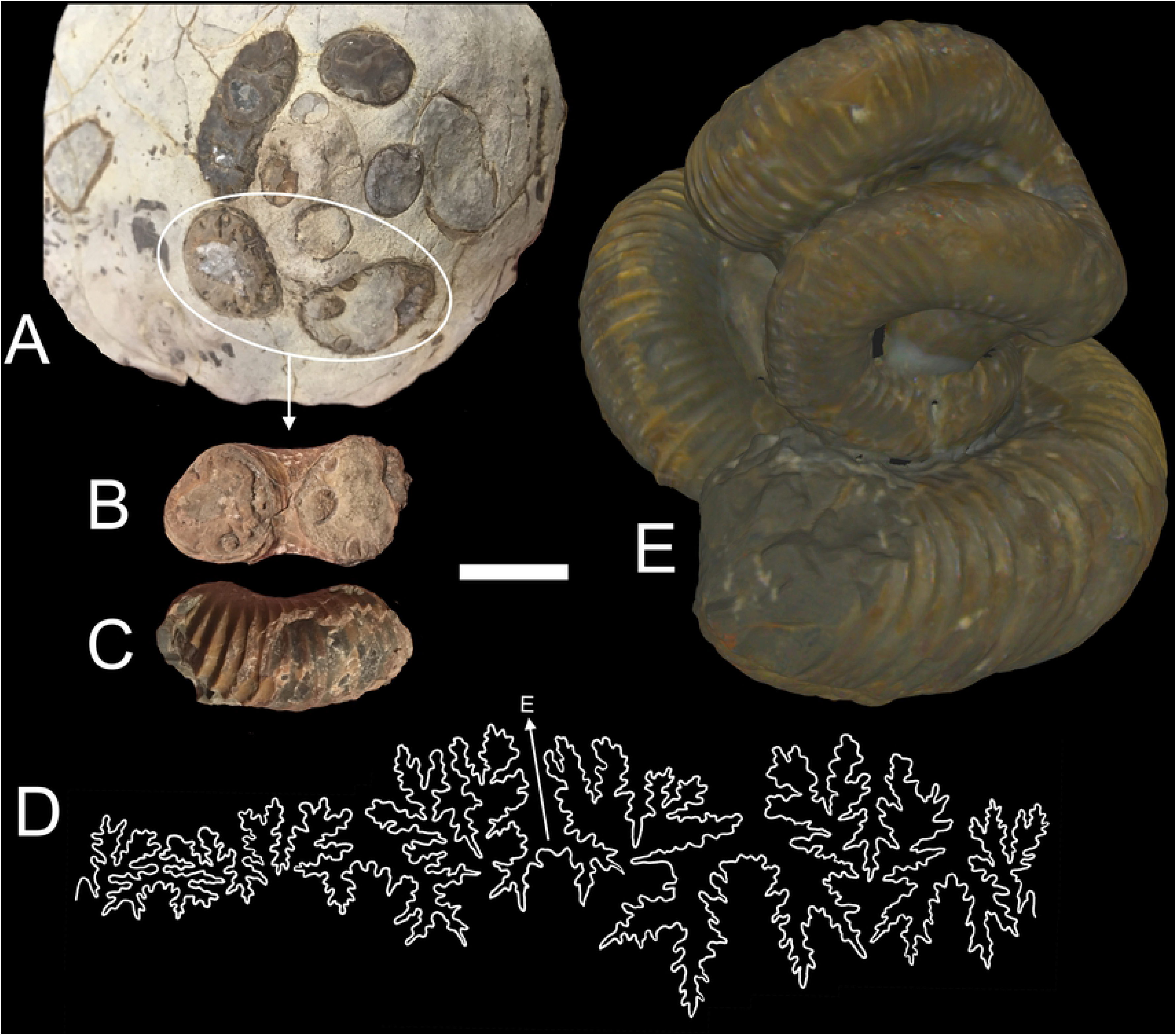
Specimens Used for Shell Reconstruction. **A**, Original concretion containing NMNS PM35490. Umbilical view (**B**) and ventral view (**C**) of the shell section used to record the suture pattern (**D**). **E**, 3D scan of the Mikasa City Museum Specimen (MCM-A0435) used to approximate the body chamber ratio. Scale bar = 2 cm.

### Virtual modeling of the soft body and camerae

The shell constrains the size and shape of other model components that influence hydrostatics. A model of the soft body was constructed by isolating the internal interface of the body chamber, and similarly, the camerae were isolated from the phragmocone of the shell. The faces of both meshes were inverted so that the normals (vectors denoting the outside) were pointing in their proper directions. The ammonoid soft body is largely unknown; however, due to phylogenetic bracketing [67], the presence of ten arms can be inferred [68–70] with a possibly reduced soft body. A soft body resembling the consensuses of Klug & Lehmann [70] and Landman et al. [71] was constructed for *Nipponites* and unified to the repaired, isolated internal body chamber mesh. The camerae were later partitioned into fractions of cameral liquid and cameral gas for hydrostatic calculations. Both cameral liquid and cameral gas were assumed be evenly distributed in the phragmocone; a reasonable assumption based on the retention of cameral liquid via the pellicle and surface tension along septal margins [72]. This yielded mass distributions of the fractions of cameral liquid and cameral gas that have the same centers as the center of volume for all camerae.

### Modeling changes in shell morphology throughout ontogeny

The final hydrostatic model of the adult *Nipponites mirabilis* was used to derive a total of 14 models representing different life stages. This was accomplished by deleting the septa in the phragmocone and deleting the adoral portion of the body chamber so that the proper body chamber ratio was maintained throughout ontogeny. The total curvilinear distance along the venter from the apex to the aperture was normalized by this same distance for the terminal stage, yielding a proxy for the age of each model (in terms of a relative percentage through the individual’s lifespan).

### Hydrostatic calculations

Neutral buoyancy occurs when the sum of organismal mass is equal to the mass of water displaced. The proportion of camerae to be emptied of cameral liquid relative to the total available cameral volume (Φ) that satisfies a neutrally buoyant condition was computed with the following equation (after Peterman et al., 2019a):

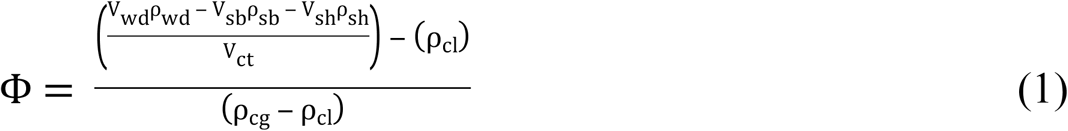

Where V_wd_ and ρ_wd_ are the volume and density of the water displaced, V_sb_ and ρ_sb_ are the volume and density of the soft body, V_sh_ and ρ_sh_ are the volume and density of the shell, ρ_cl_ is the density of cameral liquid, ρ_cg_ is the density of cameral gas, and V_ct_ is the total cameral volume of the phragmocone. A soft body density of 1.049 g/cm^3^ is preferred based on the measurement of *Nautilus* soft body by Hoffmann & Zachow [73] that was later averaged by a seawater-filled mantle cavity and thin mouthparts by Peterman et al. [38]. A shell density of 2.54 g/cm^3^ was adopted from Hoffman & Zachow [73]. The cameral liquid density of 1.025 g/cm^3^ [74] and cameral gas density of 0.001 g/cm^3^ are used in the current study.

The total center of mass is weighted according to each material of unique density (i.e., the soft body, shell, cameral liquid, and cameral gas in the current study). Each individual center of mass for the soft body, shell, cameral liquid, and cameral gas were computed in MeshLab [75] and the total center of mass was computed with the equation:

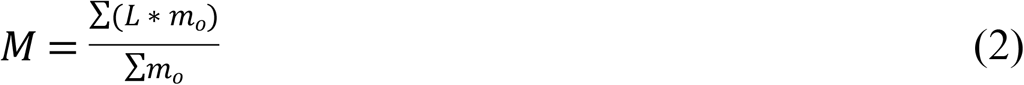

Where M is the total center of mass in a principal direction, L is the center of mass of a single object measured with respect to an arbitrary datum in each principal direction, and *m*_*o*_ is the mass of any particular object that has a unique density. Equation 2 was used in the x, y, and z directions to compute the coordinate position of the center of mass.

The center of buoyancy (B) is equal to the center of volume of the medium displaced by the external model. A model denoting the exterior interface of *Nipponites* was constructed from the external shell and soft body protruding from the aperture and its center was computed in MeshLab.

The static orientation of the total model occurs when B and M are vertically aligned. The hydrostatic stability index is computed from these centers.

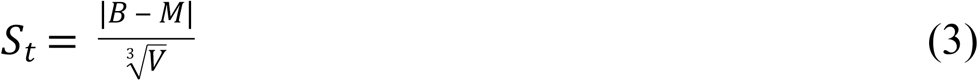

The separation between the centers of buoyancy (B) and mass (M) is normalized by the cube root of the organismal volume (V; equal to the volume of seawater displaced) in order to be applied to ectocochleates with irregular coiling [20].

Apertural angles (θ_a_) were measured with respect to the vertical (Fig 1A). That is, angles of zero correspond to a horizontally facing soft body, angles of +90° correspond to an upward facing soft body, and angles of −90° correspond to a downward facing soft body.

Thrust angles (θ_t_) were measured with respect to the horizontal (Fig 1B) between the point source of thrust and the rotational axis. Therefore, as the thrust angle approaches zero, more energy is transmitted into horizontal movement with a lower rotational component.

Rotational thrust angles (θ_tr_) were measured between the thrust vector (perpendicular to the aperture) and the rotational axis (Fig 1C). A rotational thrust angle of 90° would allow pure rotation to take place, while angles of 0° and 180° would result in translational movement.

## Results

The unknown soft body can produce errors in buoyancy calculations depending upon its total volume. By comparing the soft body used herein with a soft body that terminates at the aperture, there is only a 0.5% difference in Φ. Similarly, the mass distribution is not significantly different between either model (a 0.7 % difference in S_t_).

Because the body chamber ratio was variable on measured specimens, this ratio was manipulated by removing one septum and adding one septum to the terminal stage model with a body chamber ratio of 42%. Removing one septum increases the total body chamber ratio to 46%. This change yields a 16% increase in Φ (to 84.6%) and a 7% increase in S_t_ (to 0.0786). Adding one septum decreases the body chamber ratio to 37%. Yielding a 10% decrease in Φ (to 65.7%) and an 8% decrease in S_t_ (to 0.0676). These changes suggest that small error (~10%) in body chamber ratio would not significantly alter calculations of buoyancy or the characteristics of the mass distribution. Small deviations from the ideal body chamber ratio took place (Table 2) during model construction. However, the body chamber ratio test suggests that their hydrostatic influences are minimal.

**Table 2.** Hydrostatic properties of *Nipponites mirabilis*.

| Stage | Age (%) | BC Ratio | $\Phi$ | $S_t$ | $\theta_a$ | $\theta_{ao}$ | L (mm) | $L_x$ (mm) | L norm | $L_x$ norm | $\theta_t$ | $\theta_{tr}$ |
| --- | --- | --- | --- | --- | --- | --- | --- | --- | --- | --- | --- | --- |
| (Crio)<br>1 | 0.18 | 42.7 | 97.3 | 0.099 | 69.5 | 69.0 | 11.65 | 11.43 | 1.236 | 1.212 | 11.1 | 7.7 |
| 2 | 0.23 | 38.6 | 82.6 | 0.101 | 74.3 | 49.5 | 10.59 | 10.59 | 0.902 | 0.902 | 1.0 | 117.8 |
| 3 | 0.29 | 40.1 | 83.5 | 0.094 | 13.8 | 14.5 | 10.04 | 9.43 | 0.726 | 0.682 | -20.1 | 117.1 |
| 4 | 0.33 | 39.7 | 76.4 | 0.093 | 99.5 | 99.5 | 9.58 | 8.61 | 0.616 | 0.554 | 26.0 | 98.2 |
| 5 | 0.38 | 42.8 | 77.4 | 0.069 | 2.0 | -30.3 | 13.28 | 11.19 | 0.773 | 0.651 | -32.6 | 135.5 |
| **6** | 0.41 | 42.7 | 75.5 | 0.082 | 22.9 | 24.1 | 14.61 | 12.23 | 0.787 | 0.659 | -33.1 | 139.1 |
| **7** | 0.49 | 42.6 | 81.7 | 0.070 | 24.1 | 5.0 | 16.64 | 14.32 | 0.765 | 0.658 | -30.6 | 124.5 |
| **8** | 0.55 | 41.6 | 79.8 | 0.083 | 31.5 | 22.4 | 19.51 | 16.20 | 0.821 | 0.682 | -33.9 | 143.0 |
| **9** | 0.60 | 42.2 | 79.8 | 0.083 | 31.6 | 18.8 | 18.88 | 16.49 | 0.753 | 0.658 | -29.1 | 113.4 |
| **10** | 0.71 | 41.8 | 81.3 | 0.079 | 34.5 | 15.1 | 21.92 | 21.72 | 0.758 | 0.752 | -7.6 | 123.2 |
| **11** | 0.77 | 41.2 | 76.3 | 0.075 | 17.3 | -12.7 | 22.18 | 17.38 | 0.720 | 0.564 | -38.4 | 122.0 |
| **12** | 0.83 | 43.5 | 75.4 | 0.076 | 50.1 | 39.8 | 27.54 | 27.38 | 0.835 | 0.830 | -6.2 | 154.2 |
| **13** | 0.88 | 39.1 | 72.2 | 0.072 | -8.2 | -11.1 | 27.66 | 24.86 | 0.807 | 0.726 | -26.0 | 105.9 |
| **(Term)**<br>**14** | 1.00 | 42.1 | 73.1 | 0.073 | 19.9 | 30.0 | 28.10 | 23.44 | 0.781 | 0.651 | -33.5 | 131.1 |

Hydrostatic properties computed for the 14 ontogenetic stages examined. Crio = criocone phase;
Term = terminal phase; Age% = curvilinear length for that stage normalized by the curvilinear
length of the terminal specimen; BC Ratio = curvilinear length of body chamber normalized by
the total curvilinear length at a particular stage; $\Phi$ = the proportion of the phragmocone to be
emptied of liquid for a neutrally buoyant condition; $S_t$ = hydrostatic stability index; $\theta_a$ = apertural
angle; $\theta_{ao}$ = apertural orientation if rib obliquity was ignored (normal to shell growth direction);
L = total lever arm; $L_x$ = x-component of the lever arm, norm = normalized by the cube root of
water displaced for each particular stage; $\theta_t$ = thrust angle; $\theta_{tr}$ = rotational thrust angle.

### Ontogenetic changes in hydrostatics

Hydrostatic properties were computed for 14 life stages (Figs 5 and 6; Table 2) in order to assess changes throughout the ontogeny of *Nipponites mirabilis* and other species sharing similar morphologies. *Nipponites mirabilis* has the capacity for neutral buoyancy at all life stages, retaining liquid between approximately 3% and 28% of the total cameral volumes. After the juvenile criocone phase, Φ decreases and stabilizes at its lower values (Fig 7). Hydrostatic stability (S_t_) follows a similar decreasing trend and does not significantly oscillate (Fig 7). These hydrostatic stability index values ranging between approximately 0.10 and 0.07 are sufficiently large enough to orient the living cephalopod to maintain some static orientation during all of the examined ontogenetic stages. The orientation of the aperture (θ_a_) oscillates in a complicated fashion throughout ontogeny, ranging between approximately −11 and 99 degrees (Fig 7). Apertural orientations significantly turned downwards are not observed. The juvenile criocone phase has apertural angles of about 70°, followed by complex oscillations as the alternating U-shaped bends develop. Afterwards, there is some degree of regularity in orientation, mostly exhibiting horizontal and diagonally upwards directions (Fig 7).

**Fig 5.**
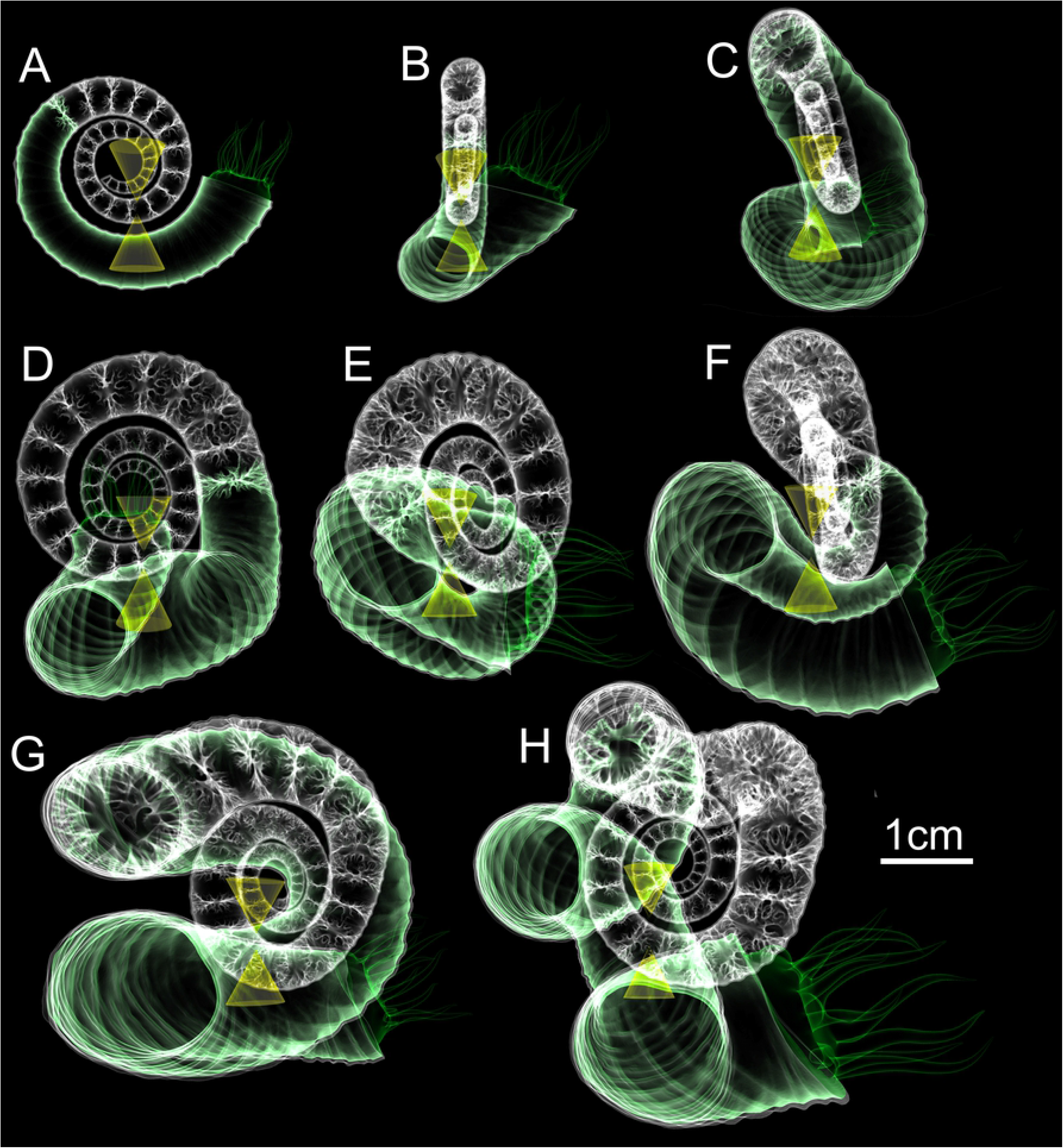
Final hydrostatic models of the first eight ontogenetic stages (A-H) of *Nipponites mirabilis*. All models are oriented so that their ventral apertures face towards the right. The tip of the upper cone corresponds to the center of buoyancy while the tip of the lower cone is the center of mass. These two centers are vertically aligned, denoting the proper static orientation assumed by living *Nipponites mirabilis*.

**Fig 6.**
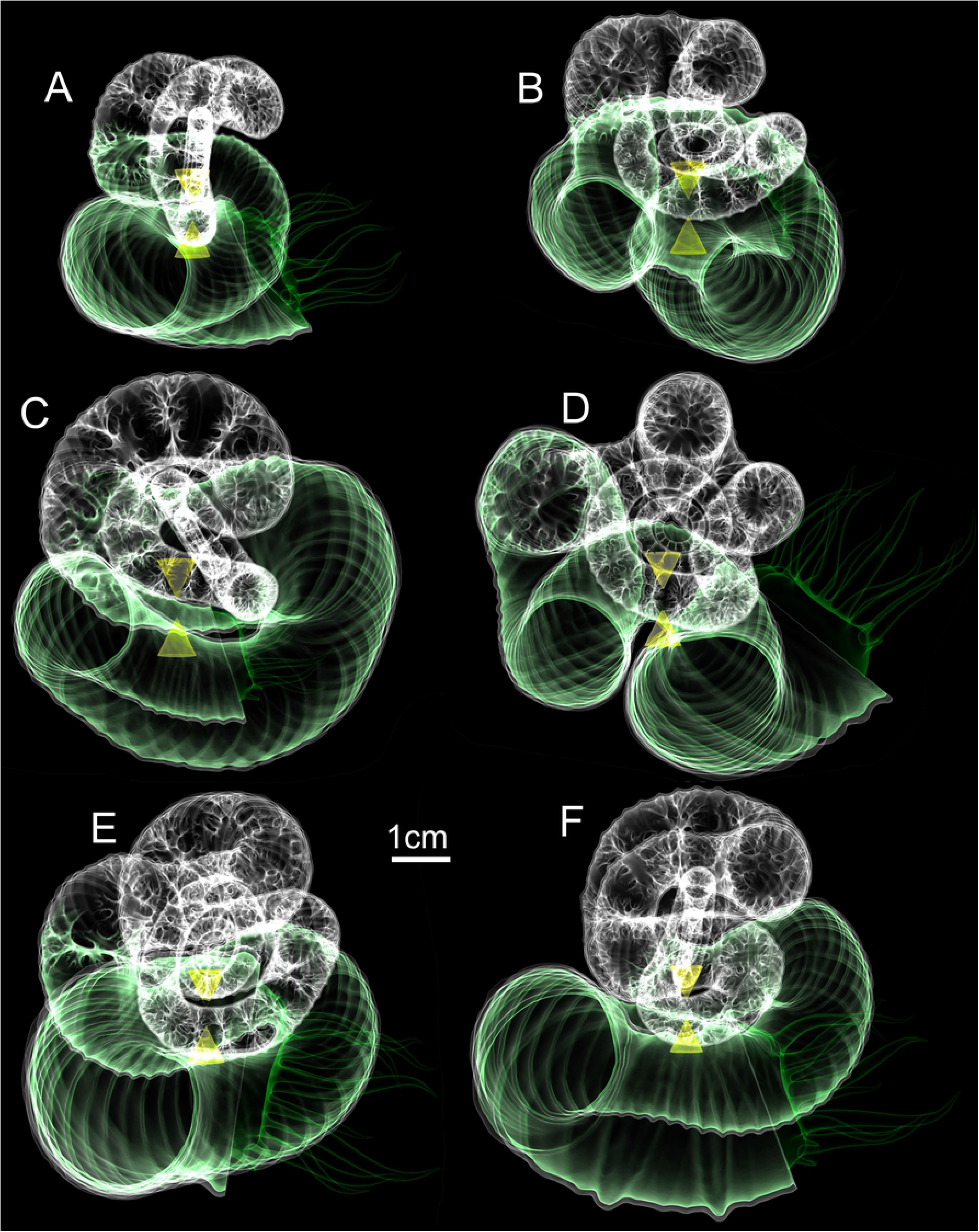
Final hydrostatic models of the last six ontogenetic stages (A-F) of *Nipponites mirabilis*. All models are oriented so that their ventral apertures face towards the right. The tip of the upper cone corresponds to the center of buoyancy while the tip of the lower cone is the center of mass. These two centers are vertically aligned, denoting the proper static orientation assumed by living *Nipponites mirabilis*.

**Fig 7.**
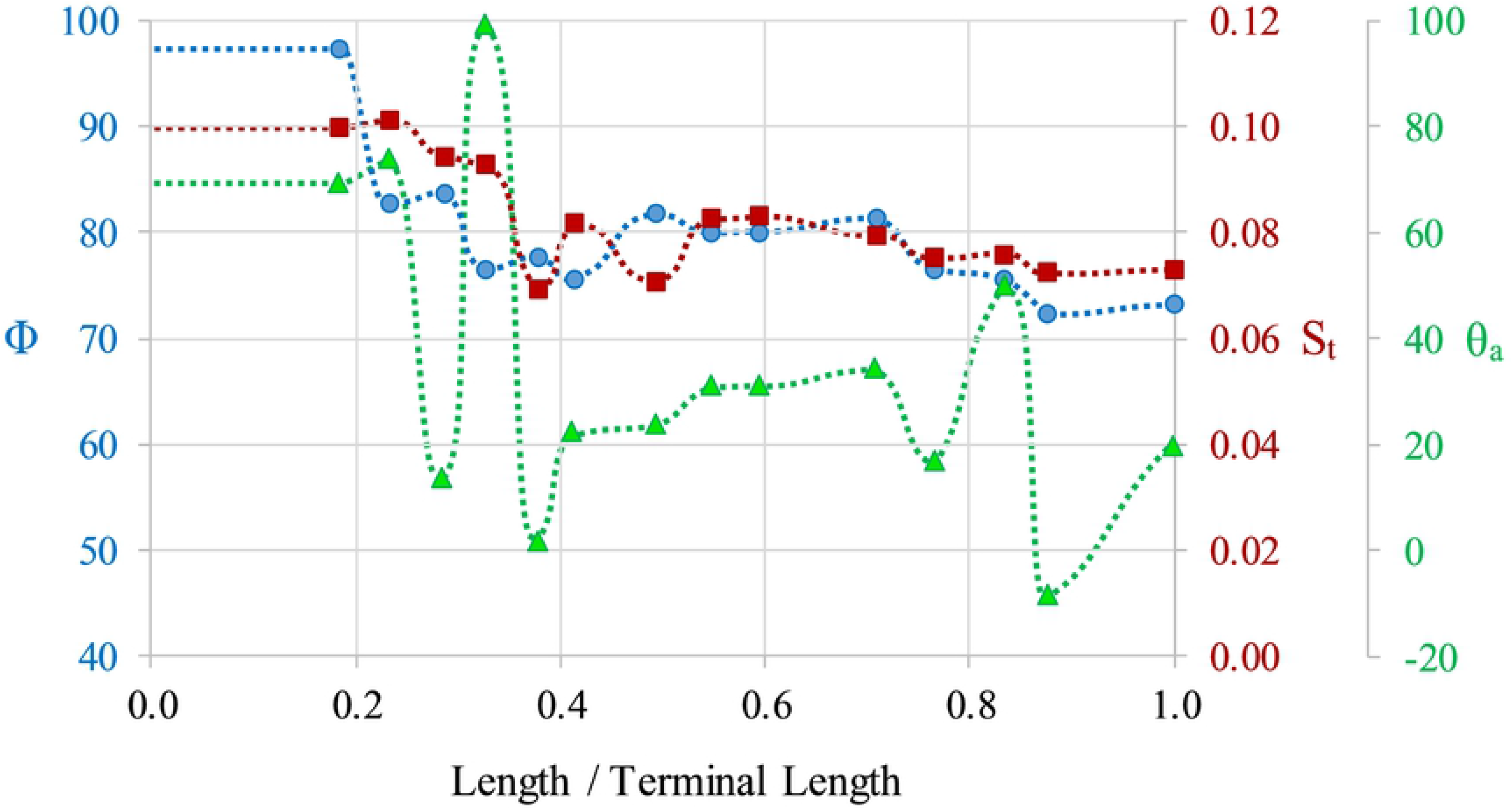
Hydrostatic Properties Computed Throughout Ontogeny. The proportion of the phragmocone to be emptied of cameral liquid for neutral buoyancy (Φ; circles), hydrostatic stability index (S_t_; squares), and apertural angles (θ_a_; triangles) as a function of age (proxied by the curvilinear length for that stage normalized by the curvilinear length of the terminal specimen). Dashed lines denote interpolations between the 14 measured stages.

### Rib obliquity and static orientation

While apertural orientations during the ontogeny of *Nipponites mirabilis* vary, horizontal to upward orientations are preferred. This is further supported by comparing the apertural angles (as denoted by the orientation of the ribs on the shell) with the same angle if ribs were not oblique (i.e., if the aperture was perfectly perpendicular to the direction of shell growth). The obliquity of the ribs generally enhances the apertural orientation by about 10° in the upwards direction (Fig 8).

**Fig 8.**
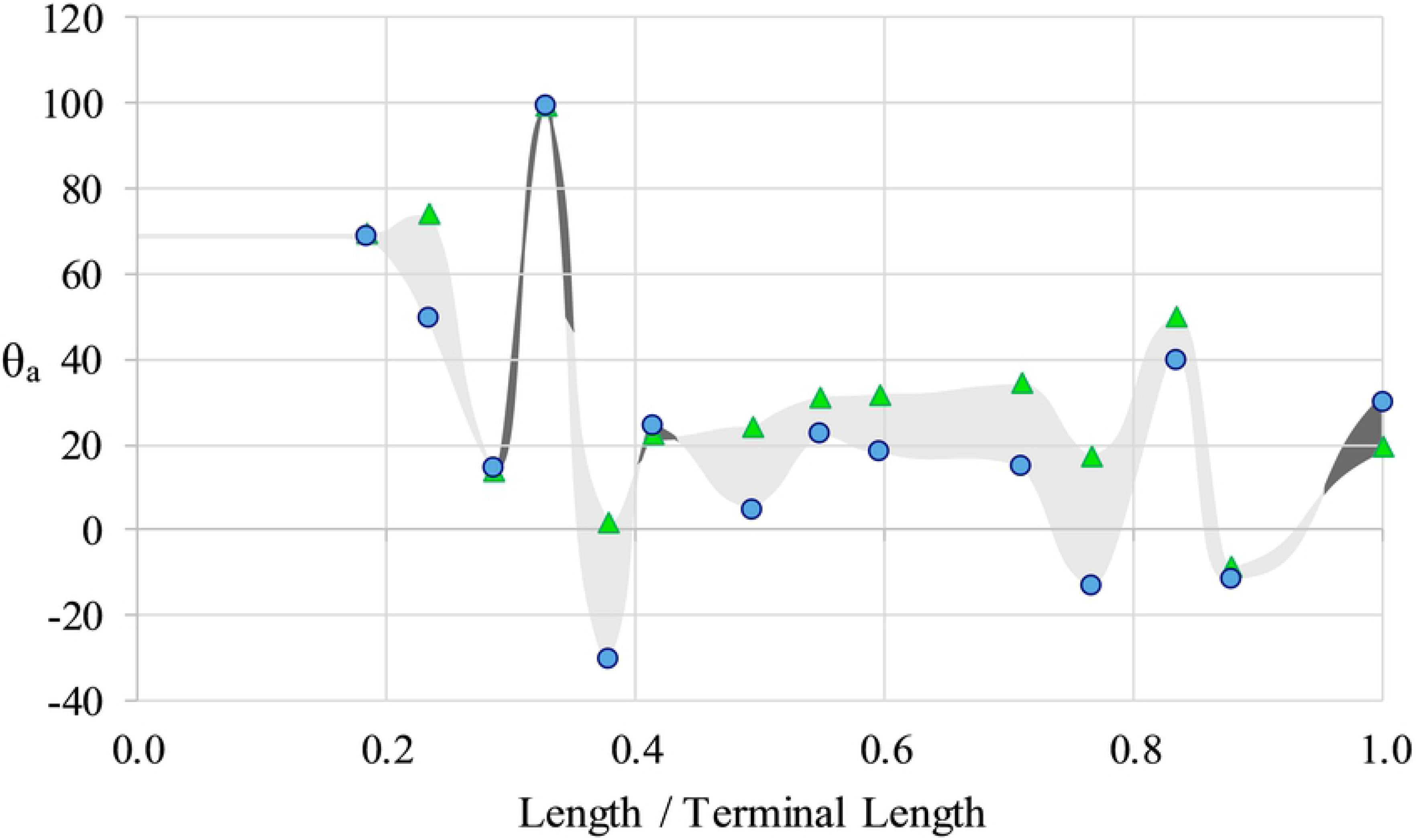
The Influence of Rib Obliquity on Orientation. Apertural angles with observed rib obliquity (θ_a_; triangles) and the angles normal to the direction of shell growth (zero obliquity; circles) as a function of age (proxied by the curvilinear length for that stage normalized by the curvilinear length of the terminal specimen). Light grey shading and dark grey shading denote rib obliquity that boosts θ_a_ in the upwards direction and downward directions, respectively.

### Directional efficiency of movement

During the juvenile crioconic phase, *Nipponites mirabilis* is well suited for horizontal backwards movement (denoted by the near zero thrust angles; θ_t_). This trend somewhat persists into later ontogenetic stages, while slightly decreasing and remaining above −40°. However, after the crioconic phase, the rotational thrust angle (θ_tr_) dramatically increases as the U-shaped bends in the shell develop; suggesting that there is a strong rotational component of movement when thrust is produced normal to the aperture (Fig 9). While the normalized lever arm lengths seem to decrease during ontogeny, sufficient torques for rotation can only be produced when the rotational thrust angle is high. Furthermore, the x-component of the normalized lever arm is not significantly lower than the total normalized lever arm during ontogeny, suggesting that the subhorizontal declination of the total lever arms would still provide significant rotational movement in ontogenetic stages after the crioconic phase (Fig 9).

**Fig 9.**
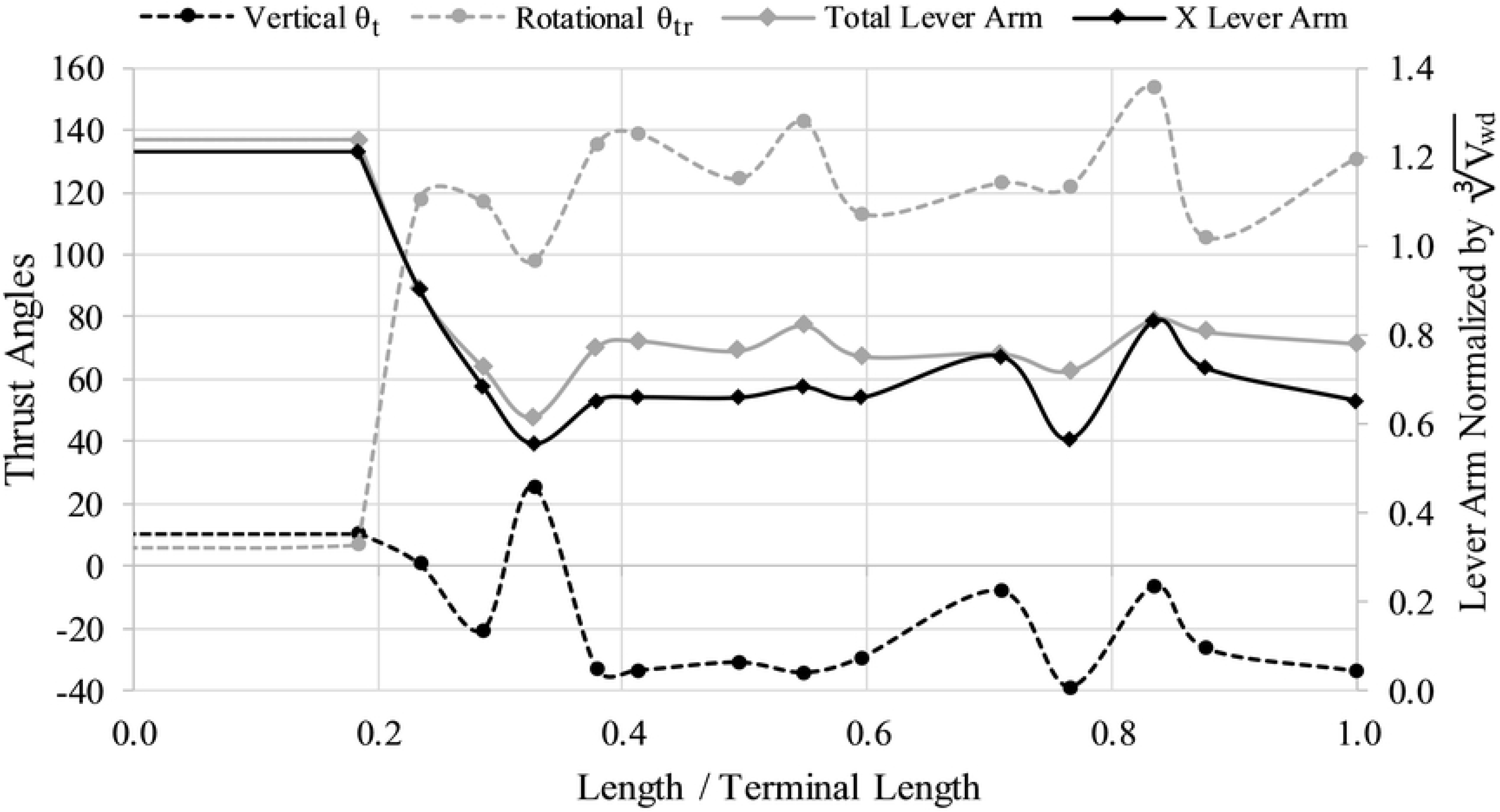
The Directional Efficiency of Movement. Thrust angles in the vertical direction (θ_t_; black dashed line), rotational thrust angles (θ_tr_; grey dashed line), and lever arms as a function of age (proxied by the curvilinear length for that stage normalized by the curvilinear length of the terminal specimen). The total lever arm (grey solid line) and x-component of that lever arm (X Lever Arm; solid black line) are both normalized by the cube root of the volume of water displaced (V_wd_) for each stage. Idealized rotation would take place with high, relative x-components of the lever arm and θ_tr_ of 90°. Idealized horizontal movement would occur with θ_t_ of 0° and θ_tr_ of 0° or 180°.

## Discussion

### The mode of life of *Nipponites*

Hydrostatic simulations reveal that *Nipponites mirabilis* had the capacity for neutral buoyancy throughout its ontogeny, retaining some amount of cameral liquid in the shell to compensate for residual buoyancy (Fig 7). These results support the buoyancy calculations of Ward and Westermann [33], who report a similar scenario for *Nipponites occidentalis*. The inferences drawn from rib obliquity most likely functioning in a neutrally buoyant setting [19] are also supported by the hydrostatic results. While the coiling of *Nipponites* is complex and somewhat resembles vermetid gastropods [23–27], considerable negative buoyancy and resultant benthic modes of life are unlikely.

Hydrostatic stability is significantly large enough for living *Nipponites* to assume static, syn vivo orientations throughout its entire ontogeny (excluding some short time after hatching when Reynolds numbers are significantly low). While the hydrostatic stability index slightly decreases throughout ontogeny, the computed values are all larger than the extant *Nautilus* (~0.05 [40]), suggesting that living *Nipponites* probably was not able to significantly modify its own apertural orientation (in terms of its vertical orientation). The highest stability in ectocochleates seem to occur for the orthocones, especially those without cameral deposits [36,40]. Lower stability values should occur for morphotypes with larger body chambers that wrap around the phragmocone (e.g., serpenticones [6,47,49]). At first glance, *Nipponites* seems to fall into this latter category at later ontogenetic stages because of the series of alternating U-bends surrounding the earlier crioconic phase and somewhat large body chamber. However, the sigmoidal soft body (which heavily influences the total mass distribution) actually seems to be somewhat confined in the vertical directions (Figs 5 and 6). That is, most of the soft body is still distributed below the phragmocone, lowering the center of mass relative to the center of buoyancy and increasing hydrostatic stability. In most cases, uncoiling of the shell seems to generally increase hydrostatic stability compared to planispiral ectocochleates [20,36,37,39,45,76].

Due to sufficient hydrostatic stability throughout ontogeny, fixed static orientations are assumed by living *Nipponites mirabilis*. That is, upward to horizontally facing orientations are preferred, while downward facing orientations were not observed in any of the examined ontogenetic stages (Fig 7). These observed orientations may have accommodated a lifestyle of feeding upon small prey in the water column, which has been proposed for other nostoceratid heteromorphs [37,45,77]. There is some period of time between about 20% and 40% of the lifespan of *Nipponites mirabilis* (after the crioconic phase and prior to the establishment of regularly alternating U-bends) where orientation oscillates between upward facing and horizontally facing. These somewhat rapid changes may have been an awkward time for these living heteromorphs. On the other hand, this irregularity infers that *Nipponites* was able to assume a functioning lifestyle regardless of these particular differences in orientation. This indifference further suggests that this heteromorph assumed a low energy lifestyle that does not demand athletic predation or predator evasion.

If the ribs of *Nipponites mirabilis* were not oblique, the static orientation of this species would be about 10° less (downward) for many of the examined ontogenetic stages (Fig 8). The obliquity of the ribs (which oscillates in magnitude throughout ontogeny [19]), therefore, assists in maintaining a generally horizontal to diagonally upward facing orientation of the soft body. Rib obliquity also suggests that the evasion of downward orientations was required to effectively function for feeding and perhaps locomotion for most stages.

### Locomotion of *Nipponites mirabilis*

The juvenile crioconic phase of *Nipponites mirabilis* would have been well suited to horizontal backwards movement with minimal rocking due to its low thrust angles and positioning of the hyponome (and thrust vector) relative to the vertical rotational axis (Fig 1A, B; Fig 9). Similar hydrostatic properties are likely for criocone morphotypes with similar proportions. Thrust angles decrease throughout ontogeny with some degree of oscillation but remain above −40°. These thrust angles at later stages suggest that significant amounts of thrust energy would still be transmitted into horizontal backwards movement, though with some degree of rocking. The subzero thrust angles post-crioconic phase result in the point of thrust located below the horizontal rotational axis suggesting that movement would be rather complicated, with oscillations in apertural angles about some horizontal axis and vertical axis, simultaneously. By examining the lever arms (normalized for each ontogenetic stage), the horizontal components of the lever arms are not much lower than the total lever arms, suggesting that rotational torque about the vertical axis during jet propulsion would be significant. After the crioconic phase, the alternating U-bends in the shell allow the thrust vector to be rotated out of alignment with the vertical rotational axis that passes through the centers of buoyancy and mass. This misalignment after the crioconic phase allows rotation about the vertical axis to take place if thrust is produced normal to the orientation of the aperture. However, this rotational thrust angle is not as ideal as torticonic (helical) heteromorphs like the turrilitids [34] and the intermediate phases of *Didymoceras* [37], which are closer to 90°. Instead, these rotational thrust angles, post-crioconic phase, fall between pure rotation (90°) and pure translation (180°) at around 135° (with some amount of variation throughout ontogeny). If the hyponome was able to bend 45° right or left, then living *Nipponites* may have been able to select between pure rotational movement and pure translational movement (influenced by some superimposition of chaotic rocking and hydrodynamic drag). This scenario depends upon the largely unknown ammonoid soft body [69,70] and propulsive mechanisms [68]. If the hyponome was not able to significantly bend, then jet thrust for post-criocone phase individuals would be transmitted into a combination of translation and rotation about the vertical axis.

The thrust angles and directional efficiency of movement provide useful information about the locomotion and feeding of living *Nipponites*. The lateral movement (and perhaps dispersal potential) of crioconic juveniles would have been on par with planispiral ammonoids (albeit with higher hydrodynamic drag), but afterwards, movement is complicated and some amount of rocking and rotation would occur. This rotational movement (pirouetting), however, could have been useful in feeding, perhaps improving the amount of space through which the living ammonoid could have searched for and captured small planktic prey. These hydrostatic properties further support a quasi-planktic, low energy mode of life for *Nipponites*.

### Complex heteromorphy in an evolutionary context

Okamoto [78] suggests that *Nipponites* originated from the nostoceratid, *Eubostrychoceras* based on comparisons of shell sculpture, early shell morphology, and stratigraphic occurrence. In a theoretical framework, the juvenile crioconic coiling of both *Eubostrychoceras japonicum* and its probable descendent, *Nipponites mirabilis*, are very similar. After this phase, the former species retains helical coiling throughout its ontogeny while the latter species alternates sinistral and dextral helical coiling [17–21,78]. While the details of the rather-sudden appearance of *Nipponites* remain unclear, the simulations of the current study infer significant differences in hydrostatic properties between these two nostoceratid genera.

*Eubostrychoceras japonicum* undergoes similar coiling patterns to the nostoceratid, *Didymoceras,* but has a longer, stretched out helical phase. Hydrostatic simulations by Peterman et al. [37] reveal that *Didymoceras* was poorly suited for lateral movement, yet adept at rotating about its vertical axis. These properties are likely analogous to *Eubostrychoceras*. While *Nipponites* has a similar ability to rotate about its vertical axis after the criocone phase, horizontal to diagonally upwards orientations are assumed instead of the likely downward diagonal orientations of *Eubostrychoceras*. For *Eubostrychoceras* to attain Nipponites-like orientations, its shell would have to coil upwards, compromising its helical coiling. Furthermore, the alternating U-bends in *Nipponites* retain some degree of lateral movement potential. Therefore, the seemingly-aberrant coiling of *Nipponites* might represent adaptations to maintaining preferred orientations and effective directions of locomotion.

The hydrostatic simulations of *Nipponites mirabilis* also provide a frame of reference for other nostoceratid heteromorphs. *N. occidentalis*, for example, exhibits a larger degree of uncoiling [33], and therefore, may have had higher stability and a larger lever arm for rotation. Similarly, throughout the late Turonian and Coniacian, a larger degree of uncoiling takes place for specimens found in successively younger strata [78]. These specimens cluster into three distinguished morphotypes [78] that may have become more stable and adept at rotation as they progressed through this time interval.

### The stigma of heteromorphy

Heteromorph ammonoids have been commonly regarded as bizarre evolutionary experiments or degenerates [8–15], and their unique coiling schemes are enigmatic in terms of their functional morphology and potential modes of life. While the inevitable phylogenetic extinction of heteromorphs (i.e., typolysis) is now rebutted [8], the stigma of this concept has persisted and is further propagated by their seemingly aberrant coiling schemes. Heteromorph ammonoids, however, were very diverse, disparate, and successful throughout the Cretaceous [45,79–81]. Furthermore, the coiling schemes of several morphotypes of heteromorph ammonoids suggest that they exploited unique solutions to manage the physical properties that constrained their modes of life by modifying their shells to serve primarily as specialized hydrostatic devices [36–39]. The hydrostatic simulations of the current study reveal that the coiling of *Nipponites*, which seems biologically absurd, does in fact confer an advantage for specific syn vivo orientations and with rotational capabilities. As suggested for several heteromorphs, the niche currently occupied by cranchid squids may be a suitable analogy for the niche once occupied by *Nipponites* [6,33,76,82].

## Conclusions

Hydrostatic analyses support a quasi-planktic mode of life for *Nipponites mirabilis* with unique forms of movement that could have enabled a planktotrophic feeding strategy. This species and other heteromorphs with similar proportions had the capacity for neutral buoyancy and were not restricted to the benthos. Throughout the ontogeny of *Nipponites*, horizontally facing to upwardly facing soft body orientations were occupied. These orientations were likely preferred for feeding on small plankton in the water column. This behavior is supported by the tendency for rib obliquity to oscillate [19], which was primarily found to upwardly adjust apertural orientations. Somewhat larger hydrostatic stability values, relative to *Nautilus*, suggest that the vertical component of the apertural orientation would not have significantly changed during locomotion or interaction with external forms of energy. A change in hydrostatics takes place between the juvenile criocone stage and the later stages consisting of alternating U-bends in the shell, specifically regarding the directional propensity for movement. Although the criocone phase of *Nipponites* likely experienced more hydrodynamic drag than planispiral ammonoids of similar size, this morphology was stable, and proficient at backwards horizontal movement. As the alternating U-bends develop, *Nipponites* is better suited for rotational movement about its vertical axis, while possibly maintaining the option to move horizontally backwards by changing the direction of its hyponome. These forms of movement were likely slow, however, suggesting that *Nipponites* assumed a low energy lifestyle while pirouetting to scan for small prey in the water column. The hydrostatic properties throughout the ontogeny of *Nipponites* contrast with those of its probable ancestor, *Eubostrychoceras* [78]. These differences in morphology along with the hydrostatic analyses in the current study infer that the seemingly convoluted coiling scheme of *Nipponites* represents unique adaptive solutions to several hydrostatic constraints, rather than random morphological aberration.

## Acknowledgements

We thank the Palaeontological Society of Japan for making CT scan data of INM-4-346 and other reference specimens available online. We also thank Keita Mori for donating NMNS PM35490. We appreciate Kei Takano for allowing us to image a private specimen used to estimate the body chamber ratio. Thanks to Daisuke Aiba, Tomoki Karasawa, and Takatoshi Tsuji for assistance with the 3D scans of MCM-A0435. Finally, we thank the Masason Foundation, the ANRI fellowship, and the JSPS (Japan Society for the Promotion of Science) KAKENHI (#18J21859) for funding a portion of this research.

---

**Table 1.** Array instructions used to reconstruct the juvenile criocone phase and the adoral portion of the terminal body chamber. These arrays were used in a piecewise manner to replicate the whorl section from the adoral direction to adapical direction by translation, rotation, and scaling in the x, y, and z directions. Asterisks denote arrays that had their origins reset to their current locations before replication. If origins were not reset, the origins of their previous arrays were used.

**Table 2.** Hydrostatic properties computed for the 14 ontogenetic stages examined. Crio = criocone phase; Term = terminal phase; Age% = curvilinear length for that stage normalized by the curvilinear length of the terminal specimen; BC Ratio = curvilinear length of body chamber normalized by the total curvilinear length at a particular stage; Φ = the proportion of the phragmocone to be emptied of liquid for a neutrally buoyant condition; S_t_ = hydrostatic stability index; θ_a_ = apertural angle; θ_ao_ = apertural orientation if rib obliquity was ignored (normal to shell growth direction); L = total lever arm; L_x_ = x-component of the lever arm, norm = normalized by the cube root of water displaced for each particular stage; θ_t_ = thrust angle; θ_tr_ = rotational thrust angle.

